# DNA origami patterning of synthetic T cell receptors reveals spatial control of the sensitivity and kinetics of signal activation

**DOI:** 10.1101/2021.03.12.434905

**Authors:** Rui Dong, Tural Aksel, Waipan Chan, Ronald N. Germain, Ronald D. Vale, Shawn M. Douglas

## Abstract

T cell receptor clustering plays a key role in triggering cell activation, but the relationship between the spatial configuration of clusters and elicitation of downstream intracellular signals remains poorly understood. We developed a DNA-origami-based system that is easily adaptable to other cellular systems and enables rich interrogation of responses to a variety of spatially defined inputs. Using a chimeric antigen receptor (CAR) T cell model system with relevance to cancer therapy, we studied signaling dynamics at single cell resolution. We found that the spatial arrangement of receptors determines the ligand density threshold for triggering and encodes the temporal kinetics of signaling activities. We also showed that signaling sensitivity of a small cluster of high-affinity ligands is enhanced when surrounded by non-stimulating low-affinity ligands. Our results suggest that cells measure spatial arrangements of ligands and translates that information into distinct signaling dynamics, and provide insights into engineering new immunotherapies.

## INTRODUCTION

Cell-surface receptors transduce extracellular signals into intracellular responses. Higher-order assemblies of signaling molecules provide important mechanisms for regulating threshold responses, amplifying signals, and suppressing noise in signal transduction (Wu, 2013). The T cell receptor (TCR) is a well-studied example of high-order assembly. Upon receptor interaction with a peptide-MHC molecule ligand (pMHC) of sufficient strength, the immunoreceptor tyrosine activation motifs (ITAMs) in the TCRζ and associated CD3 chains undergo phosphorylation, and recruit various signaling components into supramolecular microclusters 30–300 nm in diameter (Lillemeier et al., 2010; Sherman et al., 2011). Microclusters are thought to facilitate the initiation and propagation of intracellular signals by increasing the local concentration of stimulatory enzymes and excluding the signaling inhibitory molecules (Douglass and Vale, 2005; Hui and Vale, 2014; Taylor et al., 2017).

While mounting evidence has established clustering as an important mechanism for regulating TCR signaling, little is known about the nanoscale structure of TCR microclusters. Understanding how spatial arrangement of pMHC-TCR interactions within clusters affects the signaling outcome has been particularly elusive. We aim to address these problems for it provides insights about how cells convert extracellular stimuli into distinct cellular responses. Previous studies have manipulated the spatial arrangement of ligands via nanolithography (Cai et al., 2018; Deeg et al., 2013; Delcassian et al., 2013; Manz et al., 2011) or two-dimensional protein array (Ben-Sasson et al., 2021). While they offer important insights into spatial requirements of receptor-ligand engagement, their methods remain costly and technically demanding for many labs, and have significant limitations for manipulating reactions at the single-molecule nanometer scale.

To address these challenges, we sought to design a system to achieve nanoscale manipulation of ligand patterns using DNA origami, a method that relies on numerous short DNA “staple” strands to fold a multi-kilobase long “scaffold” DNA into nanoscale structures of customizable shape (Rothemund, 2006). Staple strands may also serve as handles for attaching diverse molecular components to enable novel types of biological experiments. We and others have used DNA origami to stimulate cells via patterned display of antibody fragments, peptides, or native ligands (Douglas et al., 2012; Hellmeier et al., 2021; Huang et al., 2019; Rosier et al., 2020; Shaw et al., 2019; Veneziano et al., 2020). However, relying on protein-based ligands introduces additional technical challenges, including incomplete functionalization (Hellmeier et al., 2021), and fine-tuning the strength and conformational preference of receptor-ligand interactions (Shaw et al., 2019). Previously, we developed a synthetic T cell signaling system in which we replaced extracellular pMHC-TCR interaction with DNA hybridization (Taylor et al., 2017), paving the way to develop a synthetic signaling system compatible with DNA origami structure. In this study, we combined these approaches, converting receptor-ligand interactions into DNA hybridization, which is highly tunable with DNA origami methods.

Here, we describe our results using a DNA origami signaling system compatible with live T cell imaging to investigate how specified nanometer arrangements of ligands affect MAP kinase (MAPK) signaling dynamics. For this purpose we chose an engineered receptor system (chimeric antigen receptor or CAR) that replaced the usual *αβ* TCR of the T cell with an intracellular domain derived from CD86 and the *ζ* chain immunoreceptor tyrosine activation motifs (ITAMs), to simulate the clinically important CAR-T cell systems used increasingly in cancer immunotherapy. Using this model, we found that the number and spacing of ligands within each microcluster modifies the triggering threshold and the time course of the MAPK signaling response. Specifically, ligand spacing affects the initiation time of the MAPK response and the ligand cluster size determines the duration of the MAPK signal. Low-affinity ligands, though non-stimulatory on their own, also can enhance the sensitivity of MAPK signaling when placed adjacent to stimulatory high-affinity ligands. Our results demonstrate the versatility of DNA origami in dissecting the mechanisms of transmembrane signaling and reveal the role of nanoscale ligand organization in modulating the T cell signaling response.

## RESULTS

### Design and characterization of DNA origami “pegboard” for ligand presentation

Previously, we developed a synthetic T cell signaling system in which extracellular DNA hybridization acts as the receptor-ligand interaction to trigger T cell signaling (Taylor et al., 2017). In this system, the native TCR was replaced by a DNA-based chimeric antigen receptor (DNA-CARζ) that contains an intracellular CD3ζ chain, a transmembrane domain from CD86, and an extracellular SNAP tag protein that covalently reacts with a benzyl-guanine labeled single-stranded DNA (ssDNA, “receptor strand”). DNA hybridization between receptor ssDNA with the complementary ligand ssDNA attached on the planar lipid bilayer triggers the phosphorylation of the ITAM domains of the CD3ζ and thus downstream T cell signaling.

To develop a DNA origami structure that can successfully control T cell ligand patterning, we designed a nanoscale “pegboard” that employs the ssDNA ligands as “pegs” and the DNA-CARζ as the cell-surface receptor (**Figure 1A**). The DNA origami pegboard is a square plate with dimensions of 60 × 60 nm with a height of 6 nm (**Figure 1B**). The pegboard has 72 “peg” DNA sticky ends facing the top side of the plate, which can each be extended with additional nucleotides to incorporate the ligand DNA of custom length and sequence (**Figure S1C**). The 72 “peg” staples are arranged in a 6 × 12 array with rows and columns spaced at 7 nm or 3.5 nm intervals (**Figure 1B**), respectively, which enables inter-ligand spacing from up to 38.5 nm to 3.5 nm. We also added 12 biotin staple DNAs at the bottom side of the pegboard for attachment to streptavidin-coated surfaces, and ATTO647N fluorophores at the four corners of the DNA pegboard for visualization (**Figure 1B**, **S1A**, **S1B**). Total internal reflection fluorescence (TIRF) microscopy revealed that individual DNA origami particles also displayed relatively uniform patterns with respect to size (diffraction-limited) and intensity, suggesting a uniform structure with little aggregation (**Figure 1E**, **1H**, **S2B**; variations were mainly attributable to fluorescence blinking). Photobleaching revealed four step-wise decreases in fluorescence intensity, where four consecutive bleaching events were identified (**Figure 1C**), consistent with the four-dye design. The DNA origami structure appeared to be properly folded when analyzed by agarose gel (**Figure S2D**) and negative stain TEM (**Figure 1D**, **S2E**), and had no obvious defects, deformation or aggregation.

**Figure 1.**
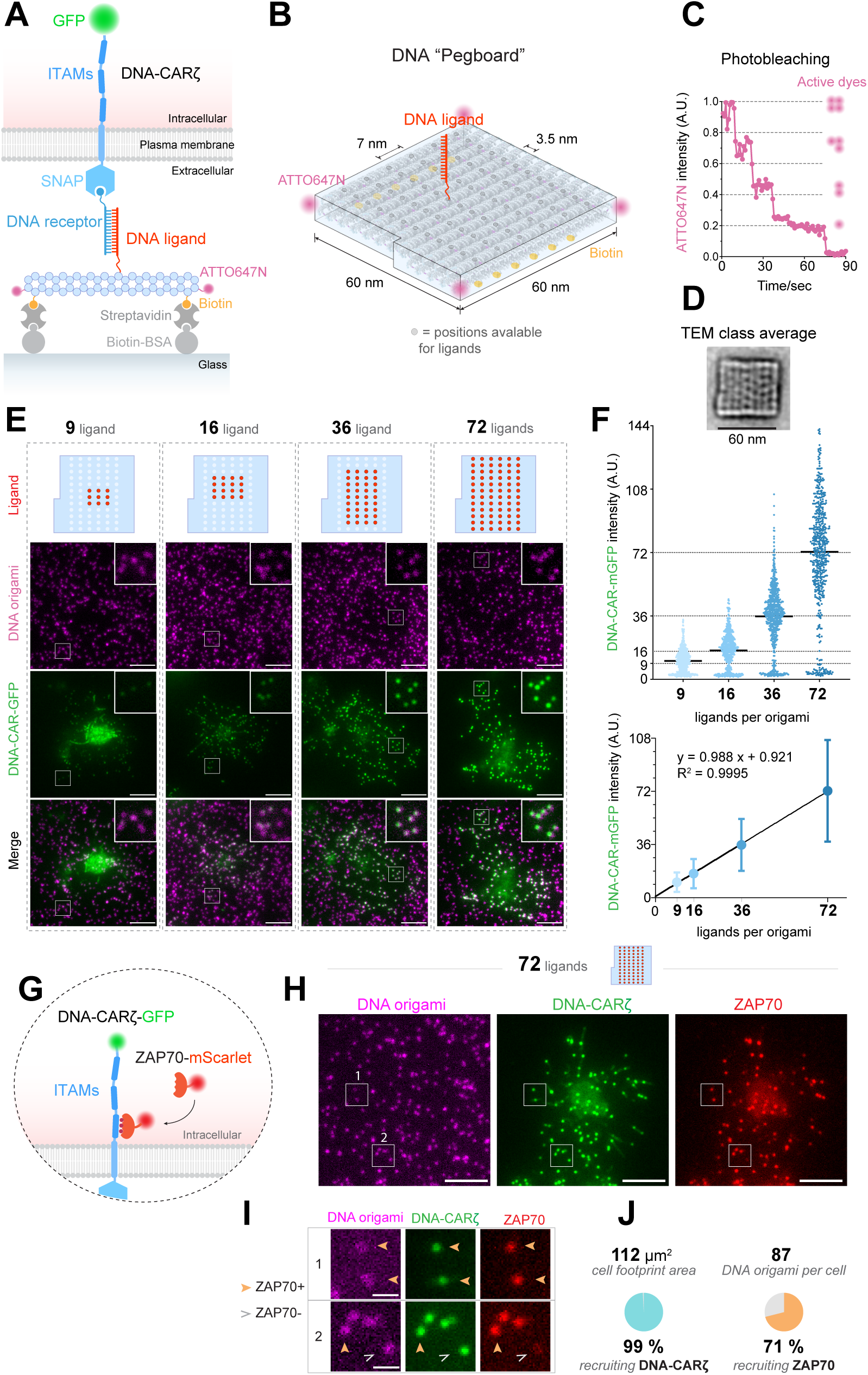
Design of the DNA origami pegboard for patterned display of ligands capable of triggering T cell signaling. (A) Schematic overview of our DNA origami-based T cell activation system. On T cell surface, the native TCR is replaced by a DNA-based chimeric antigen receptor (CAR) (DNA-CARζ) that responds to the DNA ligands (Taylor et al., 2017). The DNA-CARζ is composed of an intracellular CD3ζ chain fused with GFP, a transmembrane domain from CD86, and an extracellular SNAP tag protein that covalently reacts with a benzyl-guanine labeled single-stranded DNA (ssDNA) (“DNA receptor”). ssDNAs complementary to the receptor strands (“DNA ligands”) are displayed with customizable pattern on the top of a pegboard-shaped DNA origami architecture. DNA origami pegboard is immobilized on glass coverslip via biotin-streptavidin-biotin-BSA interactions and labeled with ATTO647N for fluorescence microscopy. (B) Designing details of the DNA origami pegboard. The DNA origami pegboard is roughly square shaped of 60 × 60 µm^2^. 72 staple DNAs on the top surface of pegboard core architecture were re-designed so that they present maximum 72 DNA ligands in a 6 × 12 array, at 7 nm or 3.5 nm intervals, respectively. The pattern and affinity of ligand-receptor interactions are by design precisely controlled via adjusting the location and sequence of the DNA ligands. Four ATTO647N dyes label the corners for fluorescence microscopy, and two rows of 6 biotin moieties were present at the bottom so that the DNA origami can mount on a glass coverslip via biotin-streptavidin interactions. (C) Fluorescence photobleaching curve for one DNA origami particle under 647 nm laser excitation. Note the five decreasing steps, which signify the bleaching events of the four ATTO647N fluorophores in succession. A.U. artificial unit (the fluorescence intensity is normalized by dividing each value by the maximal value). (D) Class averaging of the transmission electron microscopic (TEM) image of the DNA origami. Scale bar: 60 nm. (E) TIRF microscopy images of ATTO647N-labeled DNA origami and DNA-CARζ-GFP clusters in Jurkat T cells. Note that DNA-CARζ-GFP fluorescence intensity per DNA origami particle decreases as the number of ligands per DNA origami reduces. Insets show colocalization of DNA-CARζ-GFP microclusters with DNA origami particles. Scale bar: 5 µm. (F) DNA-CARζ-GFP fluorescence intensity per DNA origami particle was quantified and plotted as scattered dots (top, solid bars indicate the mean). Fitting with a linear regression model suggests that the mean DNA-CARζ-GFP fluorescence intensity is linearly proportional to the number of ligands per DNA origami (bottom, data were presented as mean ± S.D.). A.U. artificial unit (the fluorescence intensity is normalized by dividing each value by the mean value of “72-ligand” dataset, and rescaled by multiplying 72). (G-J) ZAP70 is recruited to a subset of DNA pegboard-induced receptor microclusters. (G) Schematic of the cell line used in H-J. Jurkat cells co-express DNA-CARζ-GFP and ZAP70-mScarlet. Ligand-receptor interaction triggers tyrosine phosphorylation of the ITAM domains of the DNA-CARζ-GFP, which recruits ZAP70-mScarlet. (H and I) TIRF microscopy images of a Jurkat cell stimulated by 72-ligand DNA origami particles showing ZAP70-mScarlet recruitment to a subset of DNA-CARζ-GFP microclusters. Scale bar: 5 µm. Boxed regions in H were magnified in I, showing that some DNA origami particles recruits DNA-CARζ and ZAP70 (solid yellow arrowheads), while some other DNA origami particles recruits DNA-CARζ but lacks ZAP70 (hollow grey arrowheads). (J) 1046 72-ligand DNA origami particles covered by 12 cells were examined for the percentage of recruiting DNA-CARζ and ZAP70 clustering. While the great majority (99.22 ± 1.21%) of all DNA origami particles trigger DNA-CARζ clustering, only a subset (71.04 ± 17.14%) of them recruit ZAP70.

We then examined whether the ssDNA ligand strands on the origami are capable of binding ssDNA receptor strands. DNA pegboards carrying the ligand strands recruited the receptor strands (**Figure S2A, S2B**). The amount of receptor strands recruited was linearly proportional to the number of ligands per DNA origami as intended (**Figure S2C**).

### Triggering T cell signaling using DNA origami nanostructures

Next, we examined whether the receptor DNAs on the T cell surface respond to DNA origami-carried ligands (**Figure 1E**). We used a 16-nucleotide DNA as the ligand strand that generates a high-affinity interaction (predicted off-rate > 7 hours) and shares similar linear dimensions with the native TCR-pMHC complex (Dong et al., 2019) when in complex with the SNAP tag (∼ 9.4 nm, 4 nm for SNAP, and 5.4 nm for DNA) (Taylor et al., 2017). This high affinity interaction was selected to simulate the binding of an avid antibody combining domain to its molecular target, as occurs in CAR T-cell immunotherapy (Jayaraman et al., 2020). We immobilized the DNA pegboards on glass surface via biotin-streptavidin-biotin interactions, and loaded Jurkat cells expressing DNA-CARζ-GFP to visualize receptor clustering (**Figure 1A**). The DNA pegboards recruited DNA-CARζ-GFP into sub-micron clusters (**Figure 1E**). The amount of DNA-CARζ-GFP recruited per DNA origami particle was proportional to the number of ligands per DNA pegboard (**Figure 1F**). Together, these results suggest that DNA origami can fine-tune the number of engaged ligand-receptor pairs on the T cell surface.

We then examined whether the DNA pegboard could transmit intracellular signals. TCR-pMHC interaction leads to the TCR phosphorylation, which triggers the recruitment of ZAP70, a cytosolic kinase that phosphorylates downstream targets. We tested our concept using 72-ligand DNA pegboard, which holds maximal capacity for receptor binding and thus optimizes the signal-to-noise ratio for ZAP70-mScarlet recruitment. We observed a robust recruitment of ZAP70-mScarlet to DNA-CARζ-GFP microclusters engaged by DNA origami particles (**Figure 1H–1J**). Interestingly, ZAP70 exhibits a robust initial recruitment over the first 2 min, with recruitment then decreasing to 25-50% of the peak intensity over the next ∼3 min (**Figure S3A**, **Movie S1**). The decline is unlikely due to photobleaching or special properties of mScarlet, because when we compared ZAP70-GFP and DNA-CARζ-GFP, ZAP70-GFP exhibited similar behavior whereas DNA-CARζ-GFP showed no decline in fluorescence over this time period (**Figure S3B**, **Movie S2**). DNA pegboards carrying as few as 9 ligand DNA strands also demonstrated clear recruitment of ZAP70 (**Figure S4B**). However, due to the background fluorescence of cytosolic ZAP70 (either mScarlet or GFP fusion), we could not consistently and reliably identify ZAP70 microclusters triggered by DNA pegboards carrying 6 or fewer ligand DNA strands.

TCR triggering by pMHC induces the nucleation and retrograde flow of the actin network, which carries TCR microclusters to the cell center (Murugesan et al., 2016). To examine whether ligated DNA pegboards become coupled to the actin retrograde flow as described for the native TCR, we attached DNA origami structures onto a planar supported lipid bilayer (SLB) containing biotinylated lipids, rather than fixing them onto glass (**Figure S4A**). Time-lapse images revealed that these mobile DNA origami pegboards, in complex with the DNA-CARζ microclusters, moved centripetally towards the cell center, forming the “bull’s eye” structure typical of an immunological synapse (**Figure S4C**, **Movie S3**). In summary, these results indicate that DNA pegboards are capable of triggering proximal intracellular signaling typical of the TCR.

### DNA origami pegboard revealed ERK triggering depends on ligand density

ZAP70 activation induces the recruitment of a supramolecular signalosome consisting of various signaling molecules, including LAT, Grb2 and SOS, which activates the Ras-Raf-Mek-ERK (MAPK) pathway that activates gene expression and also has been reported to act in positive feedback manner to support effective TCR signaling (Altan-Bonnet and Germain, 2005; Stefanová et al., 2003). To measure ERK activation by DNA pegboards, we engineered a new version of Jurkat T cells expressing both the DNA-CAR*ζ* and a kinase translocation reporter (KTR) that enables monitoring of ERK activities with high sensitivity in single living cells (Regot et al., 2014). Upon phosphorylation, this reporter is exported from the nucleus to the cytosol and returns to the nucleus upon dephosphorylation (**Figure 2A**). We observed ERK-KTR translocation in 33% cells stimulated with 72-ligand DNA pegboards after 12 minutes, in contrast to a background of 6% cells resting on a surface with 0-ligand DNA pegboards (**Figure 2B**).

**Figure 2.**
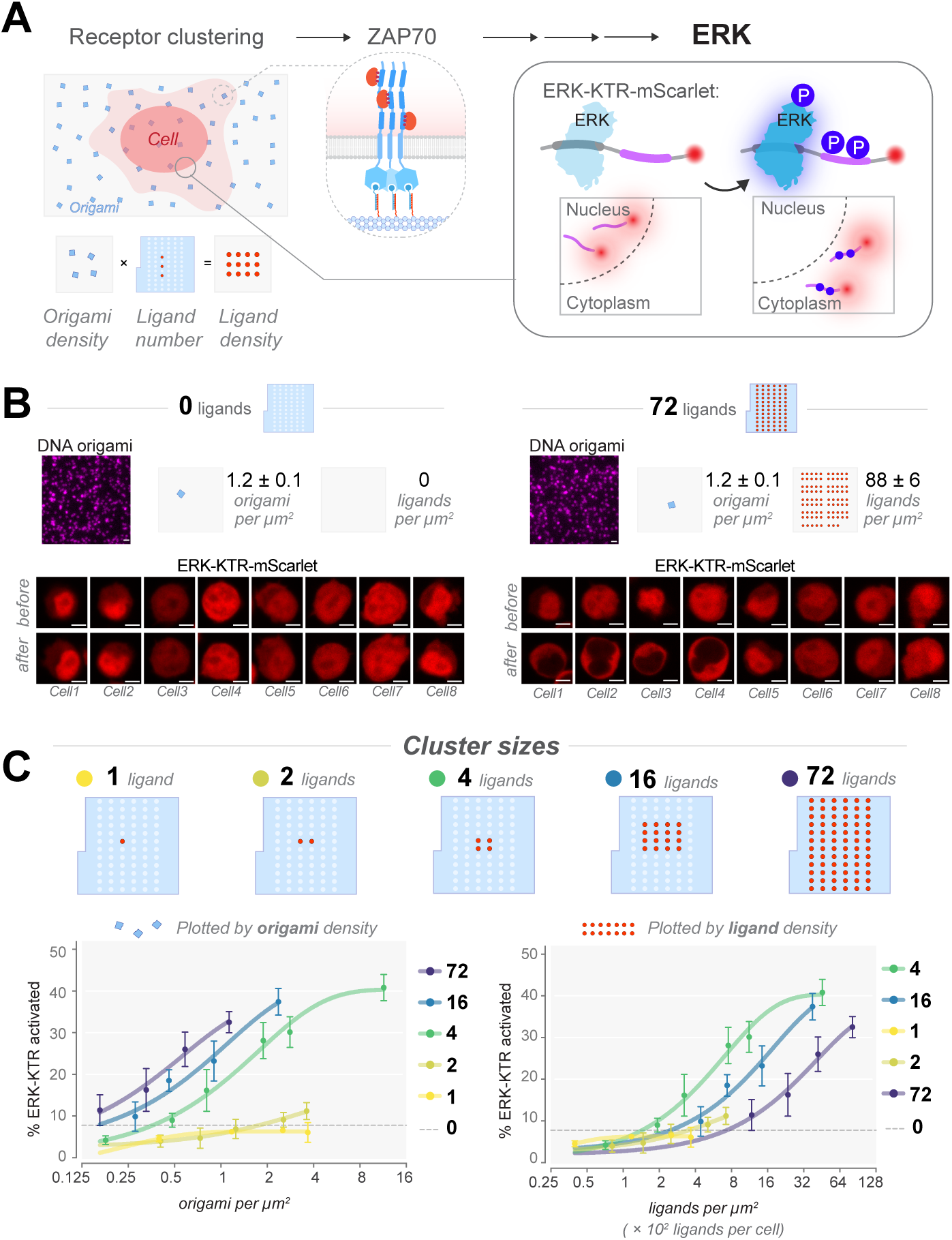
Display patterns modify the threshold ligand density to trigger ERK signaling. (A) Schematic workflow for the inspection of ERK signaling in response to DNA origami ligands displayed at a prescribed pattern. DNA origami particles were immobilized on the glass bottom of multi-well plates. DNA origami density was determined by TIRF microscopy, based on which the overall density of the ssDNA ligands was inferred (left). ERK signaling activities in cells stimulated by DNA origami ligands were monitored with the ERK-KTR-mScarlet reporter, which converts phosphorylation into a nucleus-to-cytoplasm translocation event (right). (B) Representative results from the workflow in A. DNA origami pegboards that carry either 0 or 72 DNA ligands were plated at equal density as measured by TIRF (top, scale bar: 1 µm). Jurkat cells co-expressing DNA-CARζ-GFP and ERK-KTR-mScarlet were introduced onto the DNA origami-coated glass-bottom imaging plate, and the ERK-KTR-mScarlet translocation was monitored with confocal microscopy (bottom, scale bar: 5 µm). Eight representative cells stimulated by either 0- or 72-ligand DNA origami were shown (“before”: cell landing; “after”: 18 min after cell landing). Note that 72-ligand DNA origami pegboards induced ERK-KTR translocation in a fraction of cells, whereas the 0-ligand DNA origami control did not produce such activities. (C) Impact of number of ligands per cluster on the probability of triggering ERK. DNA pegboards carrying varying ligand layouts (top) were used to stimulated Jurkat cells co-expressing DNA-CARζ and ERK-KTR. The ERK-KTR activation was monitored every 12 min over 96 min. A C:N of 2.5 was set as the threshold of ERK-KTR activation. Percentages of ERK-KTR activation were calculated at 12 min after cell landing, and plotted in terms of the DNA origami density (bottom left) or of the overall ligand density (bottom right). The estimated numbers of ligands per cell are calculated by multiplying the overall ligand density with the averaged cell footprint area (100 µm^2^). Each data point represents the mean ± SD of 3 independent replicates (> 100 cells scored per replicate). Data were pooled from two sets of experiments: one includes the 1-, 2-, and 4-ligand DNA origami, and the other include the 4-, 16- and 72-ligand DNA origami. Data for 4-ligand DNA origami from both experiments were shown, for which the results at the same DNA origami density match between two experiments and are shown as average.

Next, we sought to understand the impact of nanoscale organization of ligands on the probability of triggering ERK signaling. Specifically, we asked how many ligands per cluster were minimally required to achieve signaling competence, and whether the spatial patterning of these ligands made a difference.

To address ligand number, we designed DNA pegboards that carried various numbers of strands (**Figure 2C**, top) and compared their probabilities of triggering the ERK response within 12 min over a range of origami densities (**Figure 2C**, bottom left). 1- and 2-ligand DNA pegboards were not capable of triggering an ERK response that was significantly greater than the 0-ligand DNA pegboard control at the densities up to two origami per µm^2^, which is about 200 origami per cell. At the same DNA pegboard density (two 4-ligand DNA origami per µm^2^), the 4-ligand DNA pegboard significantly increased the probability of ERK activation (30% of cells). Similar probability of ERK activation was achieved by DNA pegboards that had higher ligand occupancy when presented at lower density (one 16- and 72-ligand DNA origami per µm^2^). In short, increased ligand occupancy per DNA pegboard increases ERK triggering sensitivity.

Next, we asked whether cells measure overall number of ligands or spatial pattern (number of clusters and/or ligands occupancy per cluster). We explored this question by replotting the dose-response curves by various DNA pegboards in terms of the overall number of ligands presented across cell contact area (**Figure 2C**, bottom right). At the same DNA ligand density (2∼8 DNA ligands per µm^2^), the 4-ligand DNA pegboard increased the probability of ERK activation compared to the 2- or 1-ligand DNA pegboards, suggesting that the ligand clustering improves ERK signaling sensitivity. As ligand occupancy per cluster further increased, they became less efficient in triggering ERK response. When the overall ligand density remained constant (∼10 ligands per µm^2^), the 72-ligand DNA pegboards triggered 10% of cells, while the 16- or 4-ligand DNA pegboards triggered 20% and 30% respectively. Overall, these results suggest that the spatial arrangement of ligands alters the threshold for T cell activation.

### Ligand clustering encodes the time course for the ERK response

The single-cell KTR reporter assay reveals information about the timing, amplitude, and cell-to-cell variation of ERK signaling (**Figure 3A**). Inspired by a growing body of evidence showing that the ERK pathway generates time-varying signals in response to different input stimuli in non-hematopoietic cells (Goglia et al., 2020; Hiratsuka et al., 2015; la Cova et al., 2017; Murphy and Blenis, 2006), we examined the time courses of ERK signals in T cells stimulated by a variety of ligand patterns.

**Figure 3.**
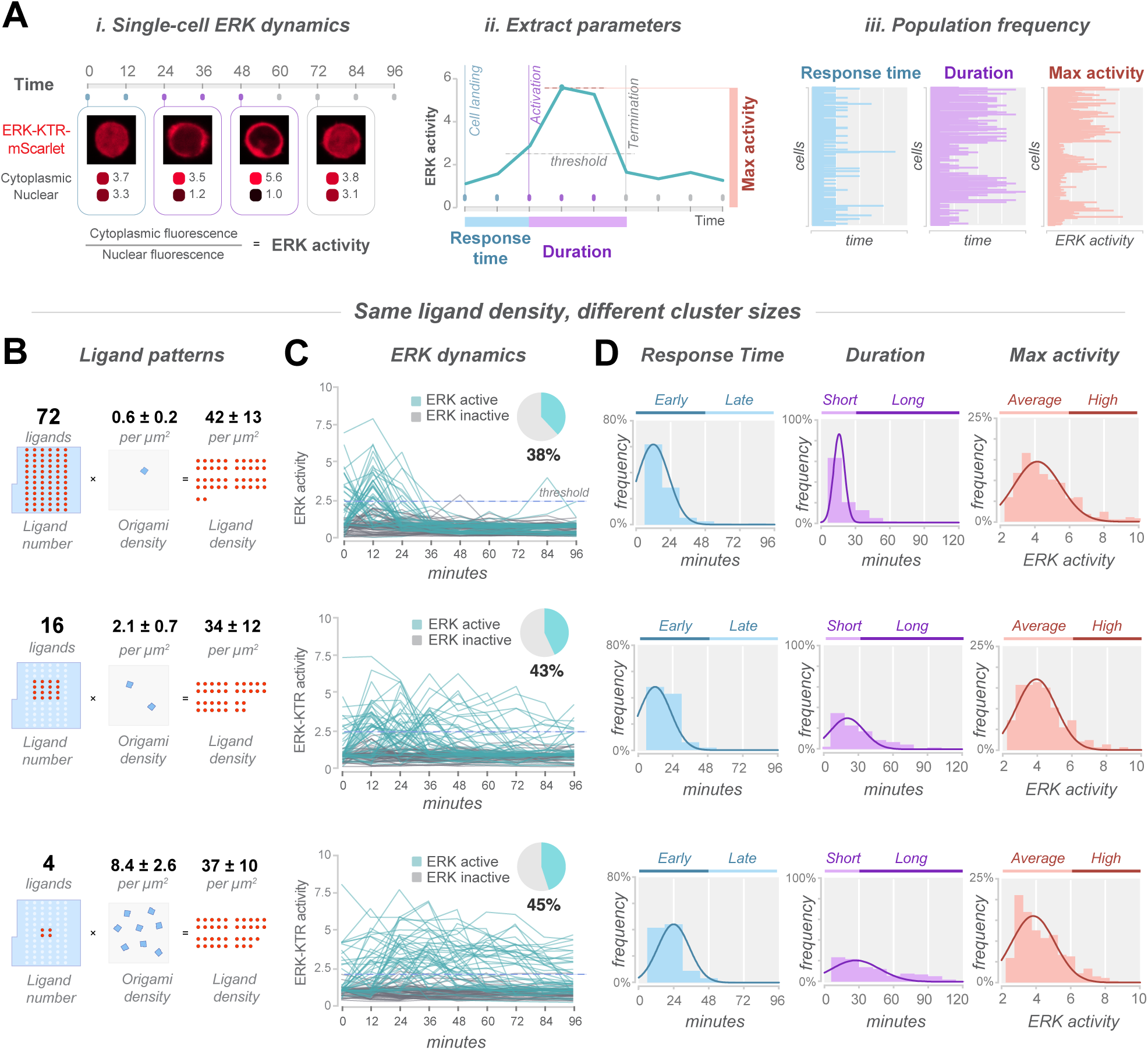
Ligand cluster size encodes the temporal course of ERK response. (A) Schematic showing the approaches of monitoring single-cell ERK activity dynamics and extracting key parameters for population-wise analysis. ERK activity was monitored at 12 min interval for 96 min (see **Figure S5** for additional notes), and calculated by the cytoplasmic-to-nuclear ratio of the ERK-KTR-mScarlet fluorescence intensity. A C:N ratio of 2.5 was set as the threshold for ERK activation. Response time is defined as the time spent from cell landing to ERK-KTR turning activated. Duration of activation is defined as the length of time ERK-KTR being positive. Maximal ERK activity was scored as the maximal C:N ratio of ERK-KTR fluorescence intensity. (B-D) Number of ligands per cluster governs the duration of ERK activation. (B) Diagrams of the DNA origamis used in C and D. DNA origami with varying numbers of ligands were immobilized on glass at respective densities so that they reached similar overall ligand densities. (C) Single-cell ERK dynamics stimulated with the DNA origami as indicated in B. ERK-KTR activity traces of 100 cells were shown for each dataset, with the ERK-active cells highlighted in cyan and ERK-inactive cells in grey. Note the sustained ERK-KTR activities in cells stimulated by the 16- or 4-ligand DNA origami, in contrast to the rapid termination of ERK-KTR activity in cells stimulated by the 72-ligand DNA origami. The pie charts indicate the percentages of ERK-active cells, which were calculated over the 96-min duration of (*n* > 600 cells pooled from 3 independent experiments). (D) Duration of activation, response time, and maximal activity scored as illustrated in Figure A. For each dataset *n* = 250 ERK-activate cells pooled from 3 independent experiments. Mean ± SD: (i) response time: “72-ligand”, 18.75 ± 11.49 min; “16-ligand”, 20.01 ± 10.63 min; “4-ligand”, 23.28 ± 13.82 min. (ii) Duration: “72-ligand”, 19.42 ± 11.32 min; “16-ligand”, 30.79 ± 20.88 min; “4-ligand”, 41.83 ± 27.88 min. (iii) max activity (C:N ratio): “72-ligand”, 4.671 ± 1.634; “16-ligand”, 4.313 ± 1.266; “4-ligand”, 4.285 ± 1.312.

Keeping the overall number of ligands presented across cell contact area the same, we varied the ligand density per origami and examined the impact of cluster size on ERK signaling (**Figure 3B**). Under the conditions of ∼34-42 ligands per μm^2^, a similar fraction (∼42%) of cells were activated by 4-, 16-, or 72-ligand DNA pegboards at varied densities over 96-min duration (**Figure 3C**). ERK signals in active cells were triggered within a similar period of time and reached a similar ERK signal magnitude (**Figure 3D**). However, remarkably, the ERK signal stimulated by the 72-ligand DNA pegboards terminated significantly earlier than the 4- and 16-ligand DNA pegboards. For example, only 10% of ERK-active cells (4% of all cells) exposed to 72 ligand origami still maintained an ERK signal at 36 min, compared to 77% of ERK-active cells (35% of all cells) exposed to the 4-ligand origami (**Figure 3C** and **3D**). In summary, our experiments reveal that ligand patterning affects the duration of the ERK response.

### Ligand spacing affects ERK signaling

We next sought to define the impact of ligand spacing on the probability of activating ERK. With a 4-ligand unit, we separated the ligands by 10 nm (“tight”) or 40 nm apart (“sparse”). We also compared these two 4-ligand origami platforms to single-ligand origamis at the same overall ligand density (**Figure 4A**, top). In an experiment in which the origami density varied, the tight 4-ligand origami was more potent than the sparse 4-ligand origami (**Figure 4A**, bottom). The sparse 4-ligand origami was only minimally more potent than the single ligand origami, when compared at a similar overall ligand density. When we examined the kinetics of ERK signaling at an equal overall ligand density (**Figure 4B**), we found that the tight 4-ligand origami has a faster ERK response time than the sparser origamis (**Figure 4C**), although the duration and the magnitude of the ERK signal were similar (**Figure 4D**). These results reveal that a denser packing of ligands leads to a faster and more efficacious ERK response.

**Figure 4.**
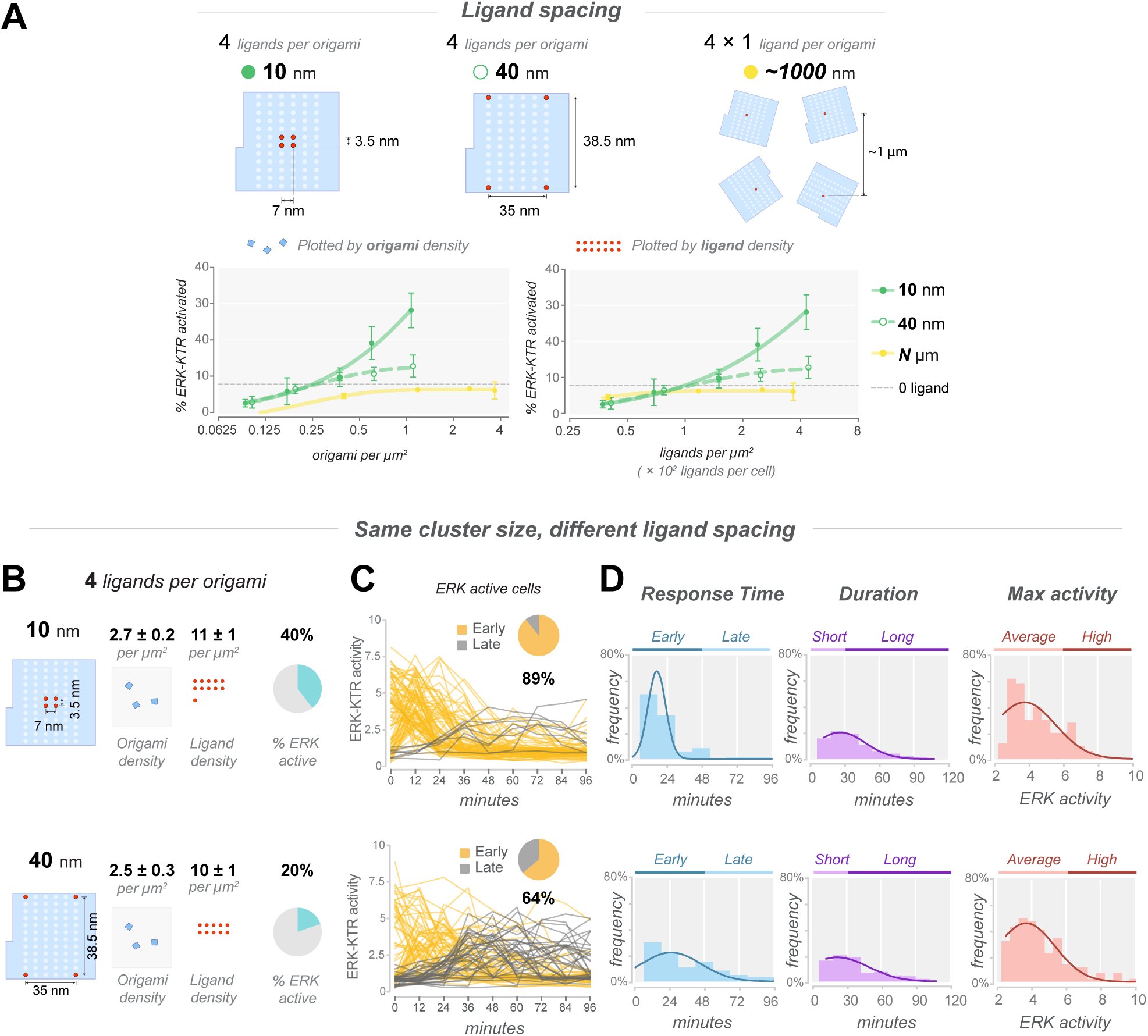
Ligand spacing modifies ERK signaling threshold and timing. (A) Impact of ligand spacing on the probability of triggering ERK. DNA pegboards carrying varying ligand layouts (top) were used to stimulated Jurkat cells co-expressing DNA-CARζ and ERK-KTR. The ERK-KTR activation was monitored every 12 min over 96 min. A C:N of 2.5 was set as the threshold of ERK-KTR activation. Percentages of ERK-KTR activation were calculated at 12 min after cell landing, and plotted in terms of the DNA origami density (bottom left) or of the overall ligand density (bottom right). The estimated numbers of ligands per cell are calculated by multiplying the overall ligand density with the averaged cell footprint area (100 µm^2^). Each data point represents the mean ± SD of 3 independent replicates (> 100 cells scored per replicate). Data were pooled from two sets of experiments: one includes the 4-ligand DNA origami with 10 nm or 40 nm spacing, the other includes the 4-ligand DNA origami with 10 m spacing and 1-ligand DNA origami with ∼1 µm spacing. Data for 4-ligand DNA origami from both experiments were shown, for which the results at the same DNA origami density match between two experiments and are shown as average. (B) Diagram of the DNA origami particles used in C and D. Both DNA origami carry 4 ligands but at different inter-ligand spacing, and were immobilized on glass at similar density. (C) Single-cell ERK-KTR activities of 100 ERK-active cells were shown for each dataset. Traces of cells that initiate ERK activation within 48 min since cell landing were colored in yellow, and those later than 48 min in grey. (D) Response time, duration of activation, and max activity of ERK-KTR activities in cells activated with the indicated DNA origami in E. Data were pooled from three independent experiments*. n* = 120 ERK-active cells out of 305 cells stimulated by the “10 nm” DNA origami, and 120 ERK-active cells out of 571 cells by the “40 nm” DNA origami. Mean ± SD: i) response time: “10 nm”, 24.06 ± 14.69 min; “40 nm”, 40.31 ± 24.27 min. ii) duration: “10 nm”, 35.40 ± 19.96 min; “40 nm”, 37.42 ± 22.43 min. iii) max activity (C:N ratio): “dense”, 4.468 ± 1.440; “sparse”, 4.475 ± 1.536.

### Closely adjacent weak ligands synergize with strong ligands to produce ERK responses

When a T cell scans the surface of an antigen presenting cell, it encounters a small number of high-affinity antigenic peptide-MHCs amidst a large number of endogenous peptide-MHCs with low affinity (Martinez and Evavold, 2015). This situation is typical for the recognition of tumor-associated antigens, which are usually expressed in extremely low levels (due to defects in their antigen processing and presentation machinery) among the vast pool of low-affinity self-peptides on the tumor cell surface (Purbhoo et al., 2006). Some studies have suggested that low-affinity pMHC ligands promote signaling by agonist pMHC (Altan-Bonnet and Germain, 2005; Krogsgaard et al., 2005; Wülfing et al., 2002), whereas other experiments observed no substantial effect (Altan-Bonnet and Germain, 2005; Spörri and Reis e Sousa, 2002).

Because most engineered CAR-T cells retain expression of an *αβ* TCR in addition to the antibody based CAR, we utilized our model system to explore if weak self-ligand interactions that might occur when such CAR-T cells encounter a tumor could facilitate signaling. We reinvestigated this question by leveraging the modularity of the DNA origami pegboard design, which allows for ligands of different affinities to be patterned in an addressable fashion. To model low affinity TCR binding, we decreased the number of hybridizing nucleotides on the ligand strand from 16 to 11, while maintaining the overall length with additional unhybridized thymine nucleotides (**Figure 5A**, top). While the half-life for dissociation of 16-mer DNA duplex is > 7 hours, the 11-mer duplex dissociates with a much faster half-life of ∼ 2 sec for the 11-mer (Taylor et al., 2017), similar to pMHC binding to a TCR (O’Donoghue et al., 2013).

**Figure 5.**
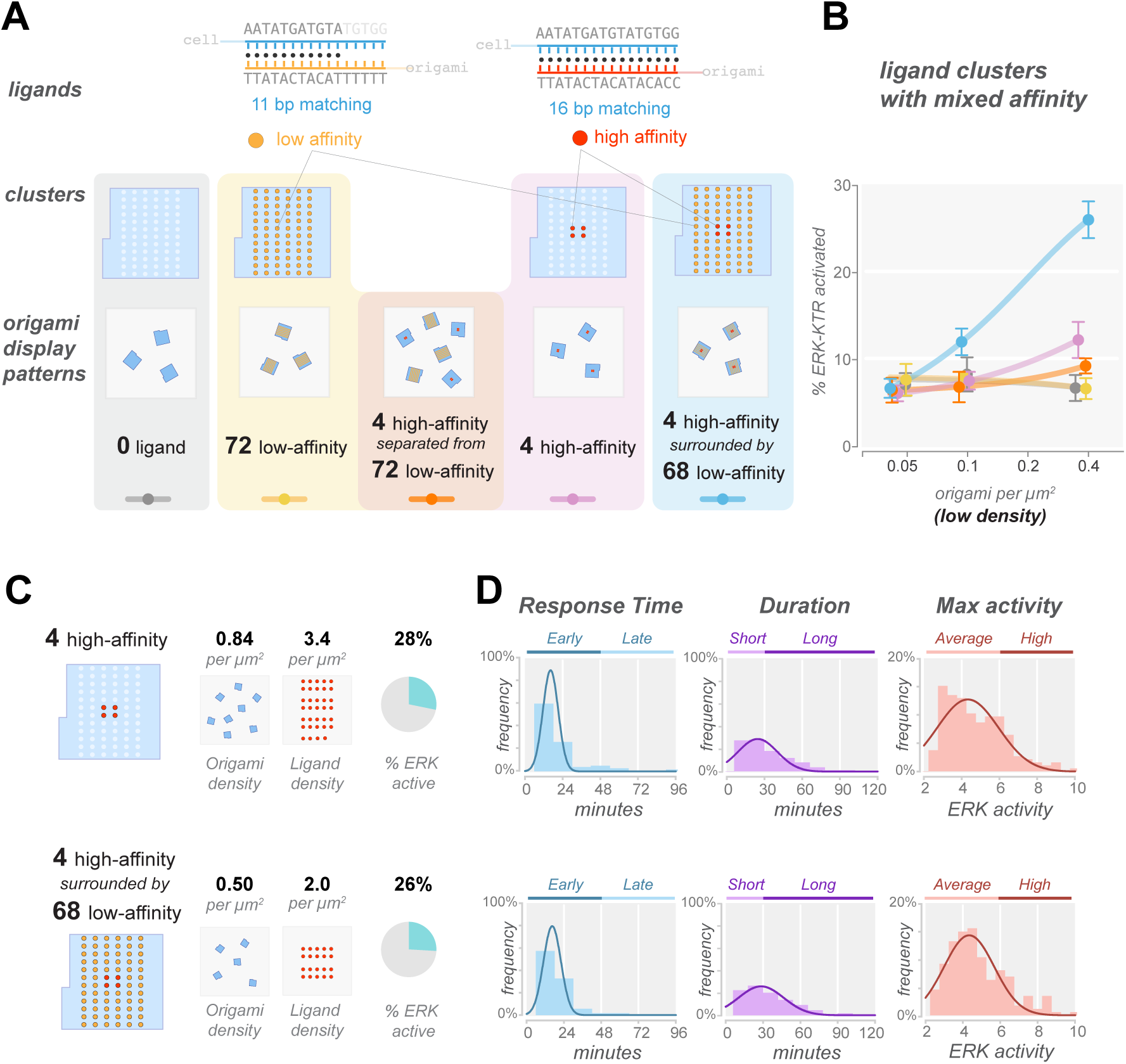
Closely adjacent weak ligands enhance sensitivity of the strong ligands. (A) Schematic of the design and display patterns of DNA origami particles used in B. (Top) as a proxy for ligands with different affinity, the numbers of hybridizing nucleotides in the ligand strands varied from 11 bp to 16 bp. (Middle) Four types of DNA origamis were designed, with the 16-mer, 11-mer, or 0-mer ligand strands patterned as illustrated. (Bottom) DNA origamis were plated on glass with the five indicated arrangements. Note that the overall density of the 16-mer ligands were the same among “4 high-affinity separated from 72 low-affinity” (orange), “4 high-affinity” (pink), and “4 high-affinity surrounded by 68 low-affinity” (blue) groups. (B) Dose-response curves for the five types of DNA origami arrangements in A. Data were plotted by the overall DNA origami density, except for the “4 high-affinity separated from 72 low-affinity” dataset (orange) which was plotted by the density of 4 high-affinity ligand DNA origami only. Data are shown as mean ± SD of 3 independent replicates (> 100 cells scored per replicate for each data point). (C-D) Additional low-affinity ligand at close adjacency to the high-affinity ligand does not modify the time course or the amplitude of ERK activities. (C) Schematics of the design and displaying density of the DNA origami particles used for D. The “4 high-affinity” or the “4 high-affinity surrounded by 68 low-affinity” DNA origami particles were plated at the specified density so that they reached a similar probability of ERK activation (data shown as mean ± SD of 3 independent replicates). (D) Frequency distribution of the duration, response time, and maximal activity of ERK stimulated by the DNA origami illustrated in C. *n* > 170 ERK-active cells out of > 700 cells pooled from 3 independent experiments were included in the analyses. Mean ± SD: (i) response time: “4 high-affinity”, 21.68 ± 18.12 min; “4 high-affinity surrounded by 68 low-affinity”, 19.18 ± 11.02 min. (ii) duration: “4 high-affinity”, 32.10 ± 19.30 min; “4 high-affinity surrounded by 68 low-affinity”, 34.97 ± 20.47 min. (iii) max activity (C:N ratio): “4 high-affinity”, 4.732 ± 1.564; “4 high-affinity surrounded by 68 low-affinity”, 4.815 ± 1.548.

To test the effects of additional low-affinity ligands, we created a DNA pegboard on which 4 high-affinity ligand strands (16-mer) were placed in the center, surrounded by 68 low-affinity ligand strands (11-mer) (**Figure 5A**, blue box). To directly test the adjacency effect and rule out the possibility that additional low-affinity ligands may collectively increase the signaling by adhesion or other means, we included a condition in which the high-affinity ligands were spatially distant from the low-affinity ligands on separate DNA pegboards (**Figure 5A**, orange box). We examined the probability of activating ERK with these ligand arrangements at extremely low ligand density, when neither the 4-ligand high-affinity DNA pegboard nor the 72-ligand low-affinity DNA pegboard triggers ERK (**Figure 2C**). The results reveal that the presence of 68 low-affinity ligands that surround the 4 high-affinity ligands heightened the probability of ERK activation (**Figure 5B**). An elevated ERK response was not observed when high- and low-affinity ligands were placed on separate DNA pegboards, suggesting that the synergistic effect requires proximity on a nanometer scale. We also compared the kinetics of ERK signaling in cells stimulated with 4 high-affinity ligand clusters with or without surrounding low-affinity ligands (**Figure 5C**). We chose a higher ligand concentration where the 4 high-affinity ligand clusters trigger on its own, so that more data were collected for the active cells. We observed similar single-cell ERK dynamics with respect to the amplitude, duration, or response time of ERK response (**Figure 5D**). These results suggest that proximal low-affinity ligands sensitize the cell to stimulation by high-affinity ligands, but do not change timing or amplitude of the ERK response.

## DISCUSSION

In this study, we demonstrate that DNA origami is a powerful method for interrogating cell signaling. Compared to nanolithography or protein 2D arrays, our DNA origami-based approach offers the following advantages: (1) The DNA origami enables precise patterning over a few nanometers, within the size range of a native TCR-CD3 complex (∼7.5 nm in diameter) (Dong et al., 2019). Conversely, the smallest nanolithography feature varies from 25 nm (Delcassian et al., 2013) to 40 nm (Cai et al., 2018). (2) The programmability and modularity of DNA structures enables simple control over multivalence, affinity and spatial arrangement, which would otherwise involve complicated protein engineering via nanolithography or protein 2D arrays. Using DNA origami, we demonstrate how ligand spacing, number, and affinity affect the threshold and kinetics of the MAP kinase signaling response, as discussed below.

### Ligand nanoclustering lowers the threshold for triggering ERK signaling

Previous studies have shown that single ligand-bound DNA-CARζs, without a DNA origami scaffold, do not stably recruit ZAP70 and do not efficiently activate ERK (Taylor et al., 2017). However, when ligated DNA-CARζs form small clusters, ZAP70 is recruited, which translates into a higher probability of ERK activation (Taylor et al., 2017). It was speculated that the DNA-CARζ clusters more efficiently exclude the transmembrane CD45 phosphatase, leading to receptor phosphorylation by the Src family kinase LCK and ZAP70 recruitment. In these earlier experiments, clusters of DNA-CARζs formed naturally and the distances between receptors could not be measured or controlled. In the present study, we could control ligand spacing. Our results are generally consistent with the idea that receptor clustering lowers the threshold for signaling. For example, we found that a tight cluster of four ligands (10 nm spacing) signals more efficiently than a sparser cluster size (40 nm spacing) (**Figure 4A**). This result is generally consistent with previous experiments using nanolithographic structures by Cai et al. (2018) that ligand clustering promotes signaling, although these structures could only probe ligand distancing down to as 40 nm (Cai et al., 2018). A super-resolution microscopic study also reported that endogenous TCR clusters trigger signaling when inter-ligand spacing decreases from 17 nm to 10 nm, although this study was correlative and did not manipulate spacing (Pageon et al., 2016).

A recent study by Hellmeier et al. also examined the effect of ligand spacing on T cell signaling using DNA origami. In their study, they used either antibodies or pMHCs as ligands interacting with the native TCR instead of a CAR, and measured calcium activation for their signaling response. They also used either one or two ligands on different sized DNA origami platforms and, in the case of two ligands, varied the spacing between the ligands. Using antibodies as ligands, they reached similar conclusions to our study regarding sensitivity. Specifically, one ligand platforms were inefficient and for two ligand origamis, decreasing the spacing between ligands from 48 nm to 20 nm significantly stimulated signaling (Hellmeier et al., 2021).

Intriguingly, the Hellmeier et al. study found that that origami platforms with a single native ligand (pMHC) could signal as efficiently as platforms containing neighboring ligand, unlike the results found with antibody ligands (Hellmeier et al., 2021). Aside from the possibility that triggering by monovalent DNA origami may reflect an artifact due to transfer of peptides from the pMHC complex to MHC on the surface of T cells for representation (Ge et al., 2002), this result indicates that pMHC interacts in some fundamentally different manner with TCR than non-native ligands (DNA ligands or antibodies). The authors suggest that one pMHC might serially trigger and activate several TCRs in a catalytic manner and that these activated TCRs could organize into signal-promoting clusters. Earlier studies by Manz et al. suggest that a minimum of four pMHC are needed in a 0.5 x 0.5 mm zone to trigger calcium signaling (Manz et al., 2011), but the Hellmeier study raises the possibility that even more TCRs might be activated in such a zone. The mechanism by which pMHC might trigger a prolonged activated state of the TCR even after ligand dissociation, the lifetime of that state, and whether/how non-ligated and ligated activated TCRs might cluster remain open questions and constitute an important topics for further investigation.

### Nanoscale patterning controls the time course of signaling dynamics

A live cell readout of ERK signaling also allowed us to examine the effects of ligand patterns on signaling dynamics. First, we find that the ERK signaling is largely an all-or-none response and the amplitude of ERK signal is not affected by ligand density or spacing. These observations are consistent with a previous study showing that the ERK signaling cascade acts in a switch-like manner and converts the diverse ligand inputs into a full-magnitude intracellular signal (Altan-Bonnet and Germain, 2005). However, we find that different ligand patterns lead to distinct probabilistic and temporal dynamics of the ERK signals (**Figure 3** and **4**).

We find that the number of ligands per origami affects the duration of the ERK response (**Figure 3**). The duration of the ERK signaling has been proposed to be a key determinant of cell fate (Murphy and Blenis, 2006), such as cell proliferation (Goglia et al., 2020), differentiation (la Cova et al., 2017), and wound healing (Hiratsuka et al., 2015) in various cell types. In mature CD8+ T cells, durations of ERK correlates with specific T cell responses, where strong and transient ERK activation induces apoptosis whereas moderate but sustained ERK activation induces cell proliferation (Wang et al., 2008). ERK signal duration has also been shown to be required for thymocyte positive selection in vivo (McNeil et al., 2005). In addition to ligand concentration or type (Goglia et al., 2020; la Cova et al., 2017; Lim et al., 2015), our results suggest spatial arrangement of ligands may be another parameter that can affect the ERK signaling response. Interestingly, at a fixed overall ligand number per cell, we find that small ligand clusters (4∼16 ligands) led to a long-lasting ERK signal than larger ligand clusters (72 ligands) (**Figure 3C** and **3D**). The mechanism of this affect is not known. One possibility is a feedback mechanism involving the tyrosine phosphatase SHP-1, which becomes rapidly activated upon receptor-ligand engagement and deactivates the TCR complex by dephosphorylation (Altan-Bonnet and Germain, 2005). Large clusters may trap SHP-1, allowing it to serially dephosphorylate neighboring DNA-CARζ receptor more efficiently compared to the same number of receptors separated on separate origamis. However, other negative machineries, including the feedback loop from ERK to Raf (Ryu et al., 2016), inhibitory factors proximal to the TCR signalosome (e.g., Csk) (Simoncelli et al., 2020), and protein abundance regulation via ubiquitination (Naramura et al., 2002) or ESCRT complex (Vardhana, 2010), may be involved in the effect that we observe on ERK signal timing.

We also show that ligand distancing delayed the initiation of the ERK response (**Figure 4B**‒**4D**). This effect might be most relevant for T cell signaling in the context of their native tissue environments. Endogenous T cells move rapidly through the tissue (5-7 µm/min) (Stoll et al., 2002) and have brief (< 10 min) encounters with the antigen-presenting cells in secondary lymphoid organs (Mandl et al., 2012). The delay in the response time of ERK signaling may provide stringent control for signaling noise (Wu, 2013). pMHCs of sufficient strength that stabilize TCRs into longer lasting microclusters might trigger a rapid ERK response within the time window of engagement between a T cell and APC in vivo.

### Triggering sensitivity to strong ligands is enhanced by adjacent weak ligands

T cells deprived of self-recognition showed a loss in T cell signaling sensitivity *in vivo* (Stefanová et al., 2002), suggesting that the highly abundant endogenous pMHCs, though not stimulatory, may play an active role in signaling regulation. However, evidence for a signaling role of low affinity pMHC is conflicting. Some studies suggest that low-affinity pMHC promote signaling by agonist pMHC (Altan-Bonnet and Germain, 2005; Krogsgaard et al., 2005; Wülfing et al., 2002), whereas other experiments observed no substantial impact (Altan-Bonnet and Germain, 2005; Spörri and Reis e Sousa, 2002). One variable in these studies is the use of previously activated versus resting or naïve T cells; failure to see low affinity ligand contribution was observed with naïve cells or previously activated cells rested for prolonged periods after initial stimulation, whereas cells tested in a narrow time period after initial activation did show responses to weak ligands (Altan-Bonnet and Germain, 2005). Jurkat, a continuously proliferating T cell tumor line, may have features closer to those of the activated cycling T cells, than the resting T cells, hence the ability to sense weak ligands. These differences in T cell state aside, in these prior experiments pMHCs of different affinities were typically assembled through chemical crosslinking, a method that lacks control over the stoichiometry or spatial organization, or simple offered to antigen presenting cells.

We revisited the impact on signaling of low affinity ligands using the DNA origami-based signaling system, which also offers the advantage of exploring the affinity and spatial arrangement of ligands. We demonstrate that, although weak ligands do not signal themselves, they can change the sensitivity, but not the timing, of ERK signaling by the strong ligands present at low density. This suggests that non-triggering weak ligands, which can nonetheless accumulate at the antigen presenting cell interface (Wülfing et al., 2002), could contribute to the active signaling of T cells and enhance responses to strong ligands, especially at low densities. Additional insight into this phenomenon may have important implications for T cell immunotherapy, and future studies could explore this affect with weak and strong pMHCs bound to a DNA origami scaffold.

### Other uses of DNA origami and relevance for immunotherapies

Our study, and other recent work (Hellmeier et al., 2021), demonstrates the utility of DNA origami for probing the effects of nanoscale organization of membrane receptors on signaling, even in regimes that are lower than super-resolution light microscopy. As we showed here, different types of ligands (here shown for high- and low-affinity ligands) can be patterned on a DNA origami surface. This approach could be extended to examine the effects of combining TCR and coreceptor ligands. In addition, it is possible to control the rigidity (Zadegan et al., 2017) and curvature of DNA origami structures (Gerling et al., 2018). Such studies could probe the effects of substrate stiffness and native membrane topology, respectively, which have been suggested to have a significant impact on T cell recognition and activation (Pettmann et al., 2018).

Our findings and future studies with DNA origami may also have important ramifications for engineering biomolecules for therapeutic uses. For example, particles functionalized with agonistic antibodies have been used to drive *ex vivo* T cell expansion for adoptive cell therapy, a treatment that uses patient lymphocytes to eliminate cancer. In this method, the prevailing strategy to achieve signaling potency is to maximize the loading of agonistic antibodies on these particles. However, our results suggest that reducing ligand loading per particle could optimally tune the duration of activation of T cells to promote sustained cell survival and proliferation, and avoid a strong but transient activation of ERK associated with apoptosis (Wang et al., 2008). Moreover, the spacing of agonistic antibodies on these particles also can be optimized to increase the probability of eliciting cellular signaling. We expect the methodology developed here would provide the basis for rapidly prototyping and optimizing nanoparticle reagents for immunotherapy.

---

**Supplementary Figure 1.**
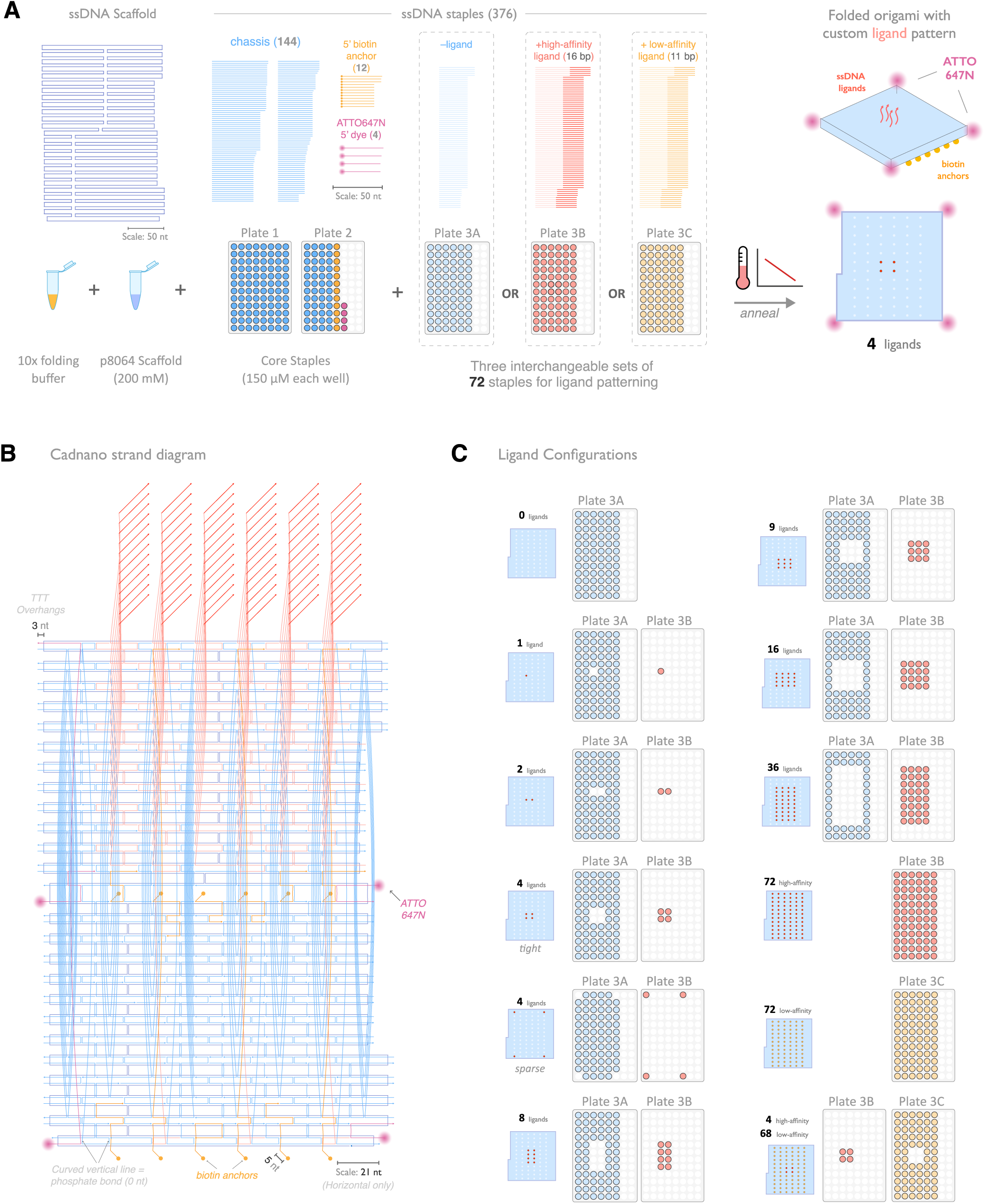
Design and production flow of DNA origami pegboard with prescribed ligand patterning. (A) 2D schematic of single-strand DNA scaffold and staples. The staple DNAs are categorized based on its location: 144 “core staples” that form the board-shape architecture, 12 biotin- and 4 fluorophore-conjugated staples that decorate the bottom and corners of the pegboard respectively, and 72 “peg” staples on the top for ligand patterning. “Peg” staples carrying varied number of base-paring nucleotides act as ligands with different affinities (See Materials and Methods for details). (B) Cadnano design strand diagram schematics. (C) To streamline the customization of ligand patterning, we synthesize the 72 “peg” staples on 96-well plate with matching layouts as those on the DNA origami pegboard.

**Supplementary Figure 2.**
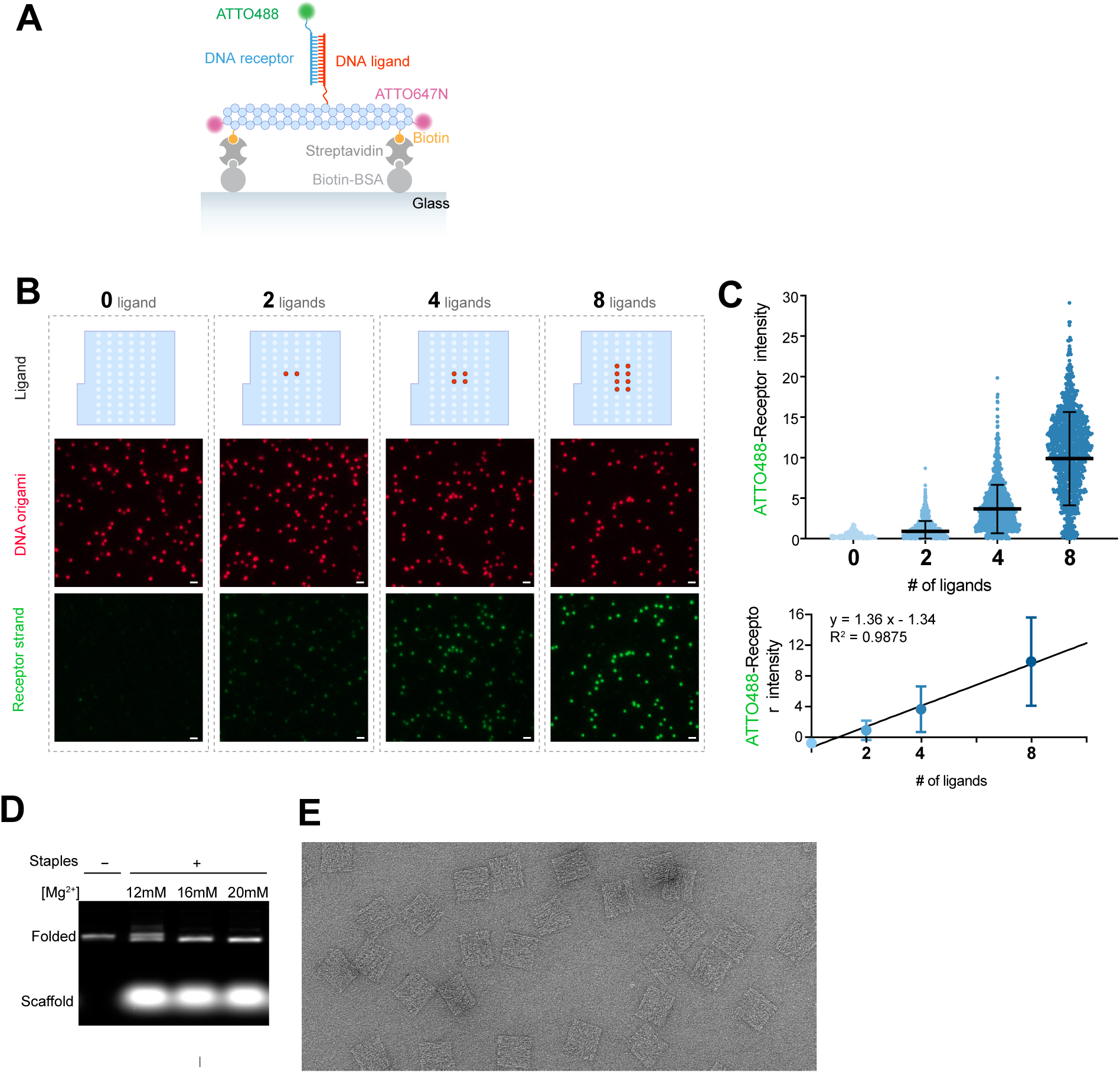
DNA origami precisely controls the number of ligands. (A) Agarose gel electrophoresis analysis (stained with SYBR green dye) showing the products of DNA origami folding reactions. 20 mM Mg^2+^ allows optimal folding and is thus used for all the rest folding reactions of the study. (B) Representative transmission electron microscopic (TEM) image of the DNA origami. Scale bars: 60 nm. (C-E) DNA origami precisely controls the number of ligands. (C) Schematic showing the experimental setup for D and E. DNA origami is immobilized on glass via biotin-streptavidin-biotin-BSA bridges, and imaged with TIRF microscopy. To test receptor-ligand DNA binding, the ATTO647N-labeled DNA origami is treated with ATTO488-labeled DNA receptor strand and assessed for fluorescence colocalization. (D) TIRF microscopic images of the ATTO647N-labeled DNA origami particles carrying varying numbers of ligand strands and treated with ATTO488-labeled receptor strands. Note that as the numbers of ligand per DNA origami decrease, the binding of receptor DNA decreases. Scale bar: 1 µm. (E) Fluorescence intensity of the receptor strands per DNA origami was quantified and plotted as scattered dots (bottom left). Fitting with a linear regression model suggests that the mean receptor binding is linearly proportional to the number of ligands per DNA origami (top right). Solid bars in the graphs indicate mean ± S.D. A.U. artificial unit (the fluorescence intensity is normalized by dividing each value by the mean value of “8-ligand” dataset, and rescaled by multiplying 8).

**Supplementary Figure 3.**
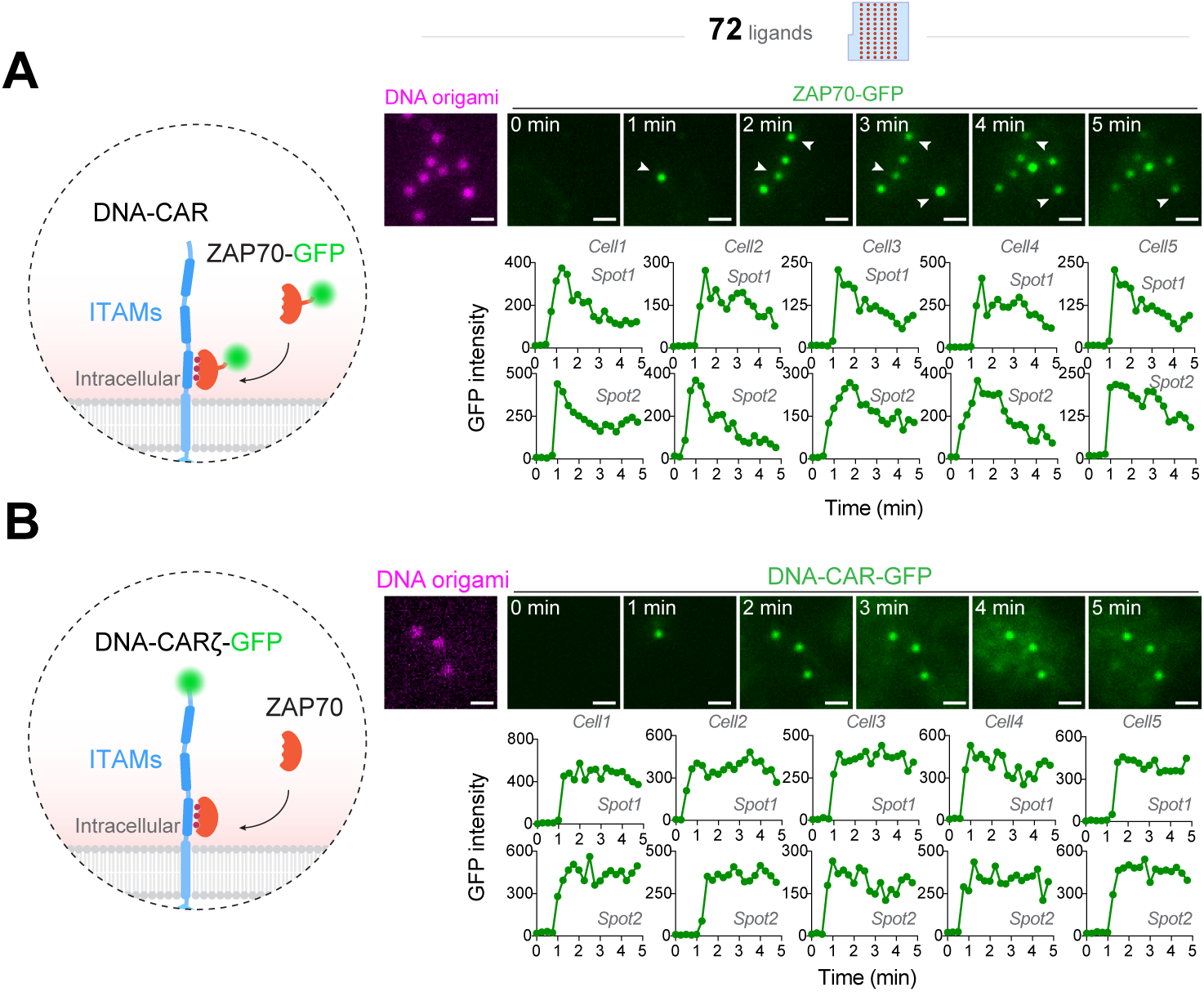
Burst-like ZAP70 recruitment to 72-ligand DNA origami. Also see Movie S1 and S2. DNA-CARζ-expressing Jurkat cells were stimulated with 72-ligand DNA origami immobilized on glass, and imaged with TIRF microscopy. Scale bar: 1 µm. Dynamics of fluorescence intensity of ten recruitment events (from five cells, two DNA origami particles each) were quantified and plotted. (A) Jurkat T cells express ZAP70-GFP and non-fluorescent DNA-CARζ. Note the robust ZAP70-GFP signal upon initial recruitment, and the relinquishing ZAP70-GFP intensity at the following time points. (B) Jurkat T cells express DNA-CARζ-GFP. Note the steady-state DNA-CARζ-GFP intensity over time.

**Supplementary Figure 4.**
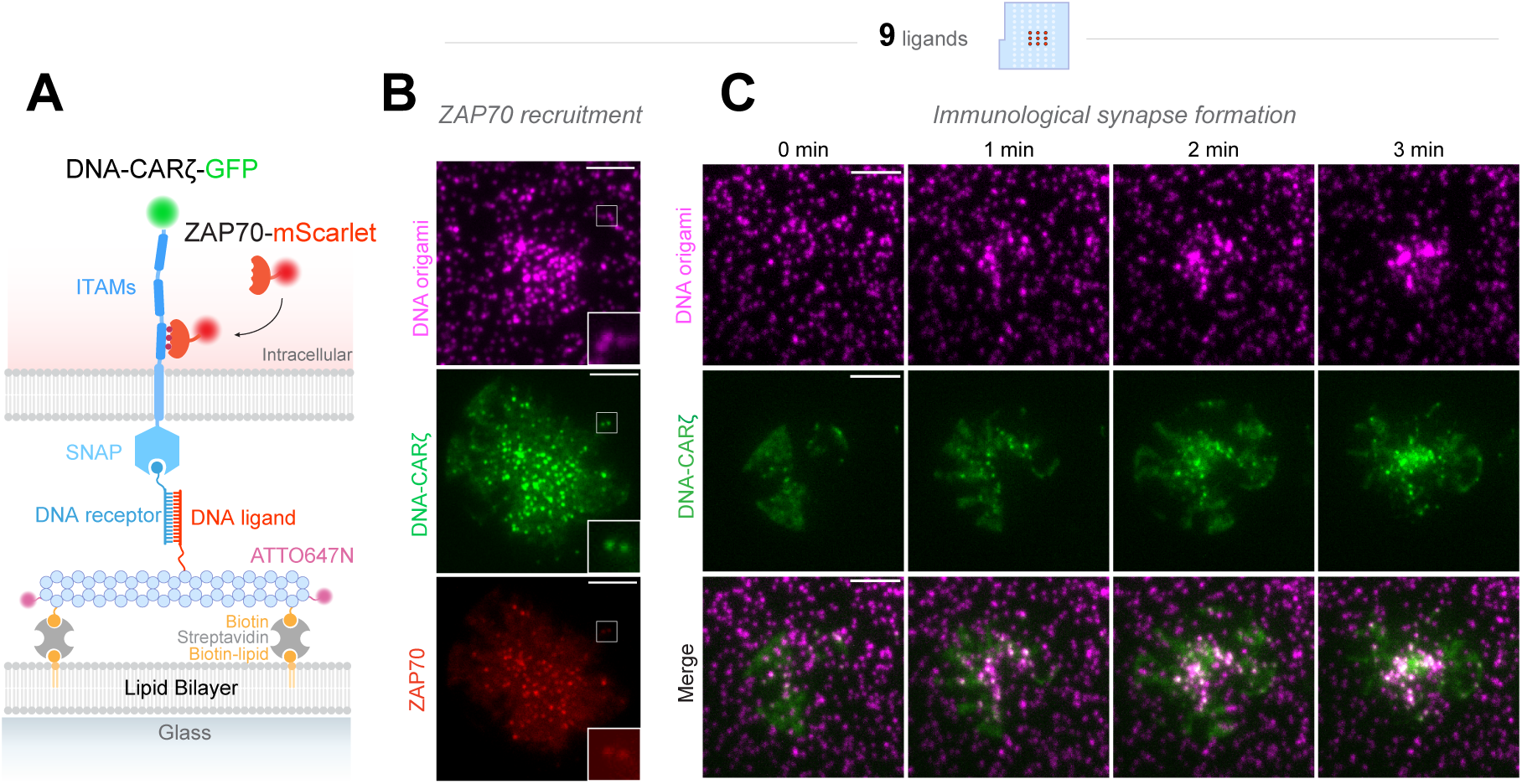
DNA origami is capable of triggering actin cytoskeleton remodeling downstream of the T cell antigen receptor. Also see Movie S3. (A) Schematic of the experimental setup. Biotin-labeled 9-ligand DNA origami particles are mobilized on the supported lipid bilayer (SLB) containing biotinylated-PE via streptavidin. Jurkat cells co-express DNA-CARζ-GFP and ZAP70-mScarlet. DNA hybridization between the ligand and receptor ssDNAs triggers tyrosine phosphorylation of the ITAM domains of the DNA-CARζ-GFP, which recruits ZAP70-mScarlet. (B) TIRF microscopy images of a Jurkat cell stimulated by 9-ligand DNA origami particles showing ZAP70-mScarlet recruitment to DNA-CARζ-GFP microclusters. Insets show the colocalization of DNA-CARζ-GFP and ZAP70-mScarlet fluorescence on individual DNA origami particles. Scale bar: 5 µm. (C) Time-lapse TIRF microscopy images of a Jurkat cell landing on SLB functionalized with 9-ligand DNA origami particles showing the formation of an immunological synapse. Note that the interactions of DNA-CARζ-GFP microclusters and DNA origami particles move centripetally and coalesce near the cell center. Insets show DNA-CARζ-GFP microclusters colocalizing with DNA origami particles. Scale bar: 5 µm.

**Supplementary Figure 5.**
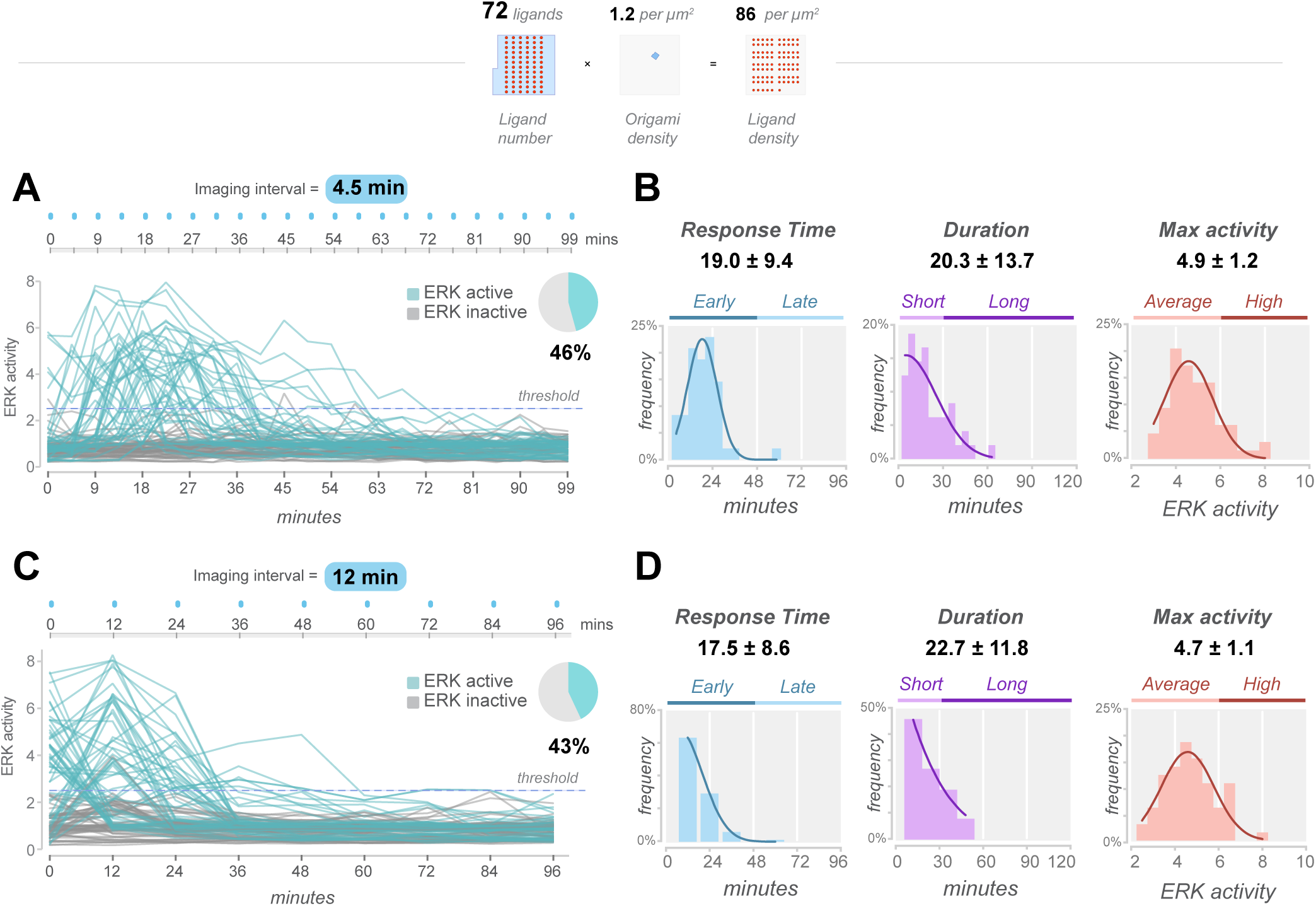
Examining ERK-KTR activities by interval imaging. 72-ligand DNA origami particles were immobilized on the glass surface at equal density, and used for stimulating Jurkat cells co-expressing DNA-CARζ-GFP and ERK-KTR-mScarlet. Single-cell ERK-KTR activity was monitored with an imaging interval of 4.5 min (A) or 12 min (B) and subject to temporal and amplitude analysis. A C:N ratio of 2.5 was set as the threshold for ERK-KTR activation. *n* = 48 ERK-active cells out of 104 cells for imaging at 4.5-min interval, and *n* = 63 ERK-active cells out of 147 cells for imaging at 12-min interval. Note that changes of the imaging intervals did not significantly shift the averaged results of the response time, duration and maximal amplitude of ERK activities. To increase imaging throughput and minimize photobleaching, all the rest single-cell ERK-KTR activity analysis in this study were carried out with a 12-min imaging interval.

## MOVIE LEGENDS

**Movie S1. ZAP70 exhibits a robust initial recruitment. See also Figure S3A.**

A Jurkat cell coexpressing DNA-CARζ and ZAP70-GFP (green) was stimulated with ATTO647-labeled 72-ligand DNA origami (magenta) immobilized on glass, and imaged with TIRF microscopy (right: merged images). Note the robust ZAP70-GFP signal upon initial recruitment over the first 2 min, and the recruitment then decreases to 25-50% of the peak intensity over the next ∼3 min.

**Movie S2. DNA-CARζ-GFP showed no decline in fluorescence over time during recruitment. See also Figure S3B.**

A Jurkat cell expressing DNA-CARζ-GFP (green) was stimulated with ATTO647-labeled 72-ligand DNA origami (magenta) immobilized on glass, and imaged with TIRF microscopy (right: merged images). Note the stable fluorescence intensity of DNA-CARζ-GFP over time, in comparison with the kinetics of ZAP70-GFP fluorescence shown in Movie S1.

**Movie S3. DNA origami is capable of triggering the formation of an immunological synapse. See also Figure S4C.**

A Jurkat cell expressing DNA-CARζ-GFP (green) was stimulated by ATTO647-labeled 9-ligand DNA origami particles (magenta) mobilized in the supported lipid bilayer (SLB), and imaged by time-lapse TIRF microscopy (right: merged images). Note that the interactions of DNA-CARζ-GFP microclusters and DNA origami particles move centripetally and coalesce near the cell center, forming the “bull’s eye” structure.

## MATERIALS AND METHODS

### DNA Origami Design

We designed the DNA origami pegboard using version 2.2 of the Cadnano software (Douglas et al., 2009). Optimization of staple routes was performed using a custom software toolkit (manuscript in preparation) developed in the Douglas lab. The structure has a two-layer design based on a honeycomb-lattice architecture (Douglas et al., 2009), which offers higher rigidity than a single-layer design. The DNA pegboard is roughly square-shaped, except for an asymmetric notch along one edge that aids in determining absolute orientation quality assessment by transmission electron microscopy.

All staple DNAs used to fold the DNA origami fall into two categories: those forming the “board” core architecture, and those forming the “peg” staples with 3’ sticky ends that terminate on the top-facing side of the board. A total of 72 “pegs” appear in a 6 × 12 array with rows and columns spaced at 7 nm or 3.5 nm intervals, respectively. These “peg” staples can be extended at the 3’ end with additional nucleotides to incorporate ssDNA sticky-ends that act as ligand strands for the DNA-CAR on the surface of the T cell. We synthesized three sets of 72 peg staples each on one 96-well plate: one set whose 3’ ends terminated flush with the pegboard surface (“–ligand”), and two sets with 3’ sticky-end extensions that contain 16 (“high-affinity”) or 11 (“low-affinity”) complementary nucleotides to the receptor strand (**Supplementary Figure 1A**).

To simplify preparation of folding stocks for a custom ligand pattern, we placed each peg staple in the well whose position on the 96-well plate matched that on the DNA origami (**Supplementary Figure 1C**).

### DNA Origami Preparation

The single-stranded DNA scaffolds used to fold the DNA origami were purchased from Tilibit (Cat No. p8064). All unmodified DNA oligonucleotides for folding the DNA origami were purchased from IDTDNA, with strands purified by standard desalting and titrated 100 µM concentration solution in water on 96-well plates. Biotin- or fluorophore-conjugated oligonucleotides were purchased separately from IDTDNA at 1 µmole synthesis scale.

DNA origami structures were folded by mixing the custom staple DNAs and scaffold DNA at a final concentration of 200 nM and 20 nM, respectively, in the folding buffer (5 mM Tris pH 8.0, 1mM EDTA, 20 mM MgCl_2_), and subjecting them to a thermal denaturation (65°C, 15 min) followed by re-annealing process (60°C to 20°C, a decrease of 1°C per 1 hour).

Folded DNA origami products were analyzed using 2% agarose gel electrophoresis in Tris-borate-EDTA (45 mM Tris-borate and 1 mM EDTA) supplemented with 11 mM MgCl_2_ and SYBR Safe. Upon sample loading, gels were run for 2 h at 80 V and subsequently scanned using a Typhoon FLA imager.

Folded DNA origami structures were purified through PEG precipitation to remove excess oligonucleotides. Products from the folding reactions were mixed with an equal volume of PEG precipitation buffer containing 15% PEG8000 (Fisher Scientific, Cat No. BP233), 10 mM Tris pH 8.0, 20mM MgCl_2_, 500 mM NaCl, and centrifuged at 16,000 g at room temperature for 25 min. The supernatant was then removed, and the (transparent) pellet was resuspended in the folding buffer. PEG precipitation procedure was repeated once more. The final pellet was resuspended in the folding buffer containing 20 mM MgCl_2_ and stored at 4°C.

### Transmission Electron Microscopy (TEM)

Purified origami structures were diluted to approximately 25 ng/μL prior to imaging. 5 μL of the diluted origami was applied to glow-discharged, carbon-coated, 400-mesh formvar grids (Ted Pella) for 1.5 min. The grid was then blotted dry on filter paper (Whatman). Washing and staining was performed by preparing a piece of parafilm with two 15-μL droplets of 1× folding buffer and two 15-μL droplets of 2% aqueous uranyl formate (electron microscopy sciences (EMS)) stain solution. The grid was dipped onto the first buffer droplet, blotted dry, dipped onto the next buffer droplet, blotted dry, dipped onto the first stain droplet, blotted dry, and then held onto the final stain droplet for 45 s before being blotted dry. Grids were then allowed to air-dry for 10 min prior to imaging. Electron micrographs were collected using an FEI TECNAI T12 transmission electron microscope using a using a 4k × 4k charge-coupled device camera (UltraScan 4000, Gatan) at 26 000× and 52 000× magnifications. Class averages were obtained using EMAN2 software.

### Functionalization of Supported Lipid Bilayer with DNA origami

All the following lipids were purchased from Avanti Polar Lipids: 16:0-18:1 POPC 1-palmitoyl-2-oleoyl-sn-glycero-3-phosphocholine (POPC; Cat #850457), 18:0 PEG5000 PE 1,2-distearoyl-sn-glycero-3-phosphoethanolamine-N-[methoxy(polyethylene glycol)- 5000] (ammonium salt) (PE-PEG5000; Cat #880220), 18:1 DGS-NTA (Ni+2) 1,2-dioleoyl-sn-glycero-3-[(N-(5-amino-1-carboxypentyl)iminodiacetic acid)succinyl] (nickel salt) (Ni2+-NTA-DOGS; Cat #790404), 1,2-dipalmitoyl-sn-glycero-3-phosphoethanolamine-N- (cap biotinyl) (sodium salt) (Biotin-Cap-PE; Cat #870277).

Small unilamellar vesicles (SUVs) were prepared from a mixture of 97.5% POPC, 0.5% PEG5000-PE, and 2.0% DGS-NTA-Ni (“DGS-NTA SUVs”) or 2.0% Biotin-Cap-PE (“Biotin-CapPE SUVs”). Lipids were dissolved in chloroform in glass tubes and dried under a stream of nitrogen gas followed by further drying in the vacuum for 2 hours. The dried lipid films were then hydrated with PBS pH 7.4 (invitrogen). The small unilamellar vesicles (SUVs) were produced by twenty freeze-thaw cycles (between −80°C and 37°C) and collected as the supernatant after centrifuge at 35,000 g for 45 min at 4°C. SUVs were stored at 4°C and used within 2 weeks.

The chambers to form supported lipid bilayer were assembled by mounting PDMS wells on glass coverslips. Glass coverslips (Ibidi Cat #10812) were RCA-cleaned followed by extensive washing with deionized water and dried under a stream of nitrogen gas. An 8-well mold was custom-made by laser cutting a 3 mm-thick acrylic board and gluing the holed board on top of another intact acrylic board. PDMS (Dow Corning) wells were made by preparing PDMS substrate mixtures according to the manufacturer’s instructions, casting the PDMS substrate mixtures into the acrylic mold, and curing at 37°C overnight. To assemble the glass-bottom PDMS cell chamber, PDMS wells were removed from the acrylic mold, cleaned together with the glass coverslips with plasma in a Harrick Plasma cleaner, and mounted on top of the glass coverslips.

To build supported lipid bilayer, 10% DGS-NTA SUV (v/v) and 1% Biotin-CapPE SUV (v/v) were mixed in PBS, deposited into each PDMS well, and incubated at room temperature at least for 1 hour. Excessive SUVs were then washed away by extraction-adding cycles (while keeping the bilayer fully submerged in the aqueous buffer).

To functionalize supported lipid bilayer with DNA origami, wells were incubated first with streptavidin (final concentration xxx, ThermoFisher Scientific Cat #S888) in PBS pH 7.4 for 10 min. The streptavidin solution was then washed out and replaced with 1x PBS supplemented with 10 mM MgCl_2_, by at least 8 extraction-adding cycles. The wells were then incubated with the DNA origami at specified concentration in PBS pH 7.4 supplemented with 10 mM MgCl_2_ for 30 min. Upon activation assay, the wells were washed extensively with the imaging buffer containing 20mM HEPES pH 7.4, 1mM CaCl_2_, 135mM NaCl, 4mM KCl, 10mM glucose, supplemented with extra MgCl_2_ up to 10 mM concentration.

### Functionalization of glass surface with DNA origami

96-well glass-bottom MatriPlates were purchased from Brooks (Catalog # MGB096-1-2-LG-L), cleaned in 5% (v/v) Hellmanex III solution (Z805939-1EA; Sigma) overnight, and washed extensively with Milli-Q water afterwards. The following day, the MatriPlates were dried under a flow of nitrogen gas and covered with sealing tapes for 96-well plates (ThermoFisher, Cat # 15036) until ready for use.

To functionalize the glass surface with DNA origami, wells on the Matriplate were incubated with 150 µL of Biotin-BSA and BSA mix solution ([Biotin-BSA] = 0.2 mg/mL, [BSA] = 0.6 mg/mL) in 1x PBS at 4°C overnight. The following, wells were washed with 1x PBS to remove excess BSA and incubated with PBS pH 7.4 containing 50 µg/mL streptavidin for 10 min at room temperature. Then wells were again washed with PBS pH 7.4 supplemented with 10mM MgCl_2_, and incubated with the DNA origami in PBS pH 7.4 with 10mM MgCl_2_ for 30 min.

### Cell Culture

Regular Jurkat cells and HEK293T cells were obtained from ATCC. A stable Jurkat line expressing ERK-KTR-mScarlet and H2B-BFP was obtained from the Ronald Germain lab.

HEK293T were grown in DMEM (Gibco, Cat No. 11965-092) supplemented with 10% FBS (Atlanta Biologicals, Cat No. S11150H) and 1% PenStrep-Glutamine (Corning, Cat # 30–009 Cl). Jurkat cells were grown in RPMI (Gibco, Cat No. 11875093) supplemented with 10% FBS, 1% PenStrep-Glutamine, and 10mM HEPES pH 7.4 (Gibco, Cat No. 15630080).

### DNA Plasmids

pHR-SNAP-TM-Zeta-mGFP, pHR-ZAP70-GFP was described earlier (Taylor et al., 2017). pHR-ZAP70-mScarlet was generated by replacing the GFP in pHR-ZAP70-GFP with mScarlet via Gibson Assembly cloning. EF1a-ERK-KTR-mScarlet lentiviral expression vector was generated by Gibson Assembly cloning based on an ERK-KTR-Clover plasmid from Markus Covert lab (Addgene #59150).

### Lentivirus Production and Generation of Stable Expressing Jurkat Cell Lines

To produce lentivirus, HEK293T cells were co-transfected with a construct of interest in pHR vector, using second generation packaging plasmids pMD2.G and psPAX2 (Addgene plasmid #12259 and #12260) using Lipofectamine LTX Reagent (ThermoFisher, Cat # 15338-100). 8∼16 hour after transfection, the media on the HEK293T cells was replaced with fresh media to remove the transfection reagent. At 48 hours post transfection, the media containing lentivirus particles were harvested, spun at 1,500 g to remove cell debris, and filtered through a syringe filter with 0.45 µm pore size. To concentrate lentivirus particles, the media was incubated with ⅓ volume of Lenti-X Concentrator (Takara, Cat No. 631231) at 4°C for 1 hour, and centrifuged at 1,500 g for 45 min at 4°C. After centrifugation, the supernatant was removed, and the pellet was resuspended in RPMI media and mixed with Jurkat cells for infection. 24 hours after infection, the media containing lentivirus particles was replaced with fresh media. Cells were analyzed a minimum of 72 hours later for infection efficiency by flow cytometry.

### Preparation of Benzylguanine-Conjugated DNA Oligonucleotides

ssDNA oligonucleotides were synthesized with 5’-amine modification (5AmMC6) by IDTDNA. The SNAP ligand, benzylguanine functionalized with N-hydroxysuccinimide ester (BG-GLA-NHS), were purchased from NEB (Cat #S9151S).

To couple oligonucleotides with benzylguanine, BG-GLA-NHS was freshly reconstituted in DMSO to a final concentration of 83mM. Amine-modified oligonucleotides were diluted in 0.15 M HEPES pH 8.5 in H_2_O, to a final concentration of 2 mM. NHS coupling to amine-modified oligonucleotide was performed by thoroughly mixing the amine-modified oligonucleotides solution with NHS ester solution aforementioned, so that the molar ratio of amine:NHS was 1:50, and the final concentration of HEPES was within 50∼100mM. The reaction was left overnight at room temperature on a rotator. The reaction product was purified next day with illustra NAP-5 Columns (Cat #17085301) to remove excessive NHS ester, using H_2_O for elution. The absorbance of the purified benzylguanine conjugated oligonucleotides at 260 nm was measured with Nanodrop to determine the molar concentration. Column-purified benzylguanine conjugated oligonucleotides were further condensed with the Savant SpeedVac DNA 130 Integrated Vacuum Concentrator System, resuspended in H_2_O (final concentration 100µM), aliquoted and stored at −20°C until use.

### Labeling DNA-CAR-Expressing Cells with Benzylguanine-Labeled Receptor ssDNA

Before activation assays, cells were pelleted, washed with 1× PBS twice, and serum-deprived for 3 hours in RPMI (Invitrogen) supplemented with 1% PenStrep-Glutamine, and 10mM HEPES pH 7.4.

Up to 1 million cells were pelleted and washed and resuspended with the imaging buffer containing 20mM HEPES pH7.4, 1mM CaCl_2_, 135mM NaCl, 0.5mM MgCl_2_, 4mM KCl, 10mM glucose. The cell pellet was then resuspended with 50 µL of imaging buffer containing benzylguanine-labeled receptor DNA (final concentration 10 µM), and incubated at room temperature for 15 min. Afterwards, cells were washed twice with at least 10mL of imaging buffer to remove excess unbound benzylguanine labeled DNA and resuspended with the imaging buffer. Notably, cells were kept in buffer containing physiological Mg^2+^ level until activation, when they were deposited onto DNA origami mounted imaging chamber where 10 mM MgCl_2_ were supplemented in the imaging buffer to preserve the integrity of the DNA origami.

### Microscopy

Imaging was performed on an inverted microscope (Nikon TiE,Tokyo, Japan) equipped with a Yokogawa spinning disk confocal and TIRF combined system (Spectral Diskovery, Ontario, Canada), a Nikon 100× Plan Apo 1.49 NA oil immersion objective and four laser lines (405, 488, 561, and 640 nm), a Hamamatsu Flash 4.0, and μManager software to run the microscope and 653 capture the images. Confocal images were captured using an Andor iXon electron-multiplying charge-coupled device camera. For TIRF imaging, a polarizing filter was placed in the excitation laser path to polarize the light perpendicular to the plane of incidence. The angle of illumination was controlled with either a standard Nikon TIRF motorized positioner or a mirror moved by a motorized actuator (CMA-25CCCL; Newport). Imaging experiments involving live cells were performed within a 37°C chamber.

### ERK-KTR Imaging and Analysis

DNA origami and cells were loaded on 96-well dark-wall MatriPlate (Brooks, Cat # MGB096-1-2-LG-L). Fluorescent images were acquired with a Nikon plan apo 40x 0.75 NA air objective lens. Image acquisition was performed using MicroManager software. Each well was imaged using the Create Grid plugin in the MicroManager multidimensional acquisition GUI. The Create Grid plugin was used to automate the acquisition of the entire well. Prior to stimulation, Jurkat cells were starved in serum-free RPMI media for 4 hours. Isolated cells were plated at low density to avoid lateral propagation of ERK activity pulses to the neighboring cells observed previously (Aoki et al., 2013).

### Image Analysis

Images were analyzed using Fiji. The same brightness and contrast were applied to images within the same panels. For measuring the receptor binding of the DNA origami, fluorescent images taken from the same field of view at 488nm and 640nm channels respectively were first background-subtracted with a sliding paraboloid algorithm (rolling ball radius = 50.0 pixels). A Gaussian filter (*σ* = 1) was then applied to reduce noise. With the 647nm image, DNA origami spots were identified using the “spot intensity analysis” plug-in developed in-house by Nico Stuurman (https://imagej.net/Spot_Intensity_Analysis), with spot radius preset at 3 pixels.

## ACKNOWLEDGEMENTS

We thank Xiaolei Su, Meghan Morrisey, Nadja Kern, and Chaim Gingold for constructive suggestions on this manuscript. R.D. was supported by a Jane Coffin Childs Postdoctoral Fellowship. T.A. was supported by a Ruth L. Kirschstein NRSA Postdoctoral Fellowship (F32GM119322) and the National Science Foundation (OAC-1740212). R.V. is an investigator of the Howard Hughes Medical Institute. S.M.D. was supported by the Army Research Office (W911NF-14-1-0507) and Office of Naval Research (N00014-17-1-2627).

